# Multi-shanks SiNAPS Active Pixel Sensor CMOS probe: 1024 simultaneously recording channels for high-density intracortical brain mapping

**DOI:** 10.1101/749911

**Authors:** Fabio Boi, Nikolas Perentos, Aziliz Lecomte, Gerrit Schwesig, Stefano Zordan, Anton Sirota, Luca Berdondini, Gian Nicola Angotzi

**Author notes:** Equal contribution (Luca Berdondini, Gian Nicola Angotzi).

## Abstract

The advent of implantable active dense CMOS neural probes opened a new era for electrophysiology in neuroscience. These single shank electrode arrays, and the emerging tailored analysis tools, provide for the first time to neuroscientists the neurotechnology means to spatiotemporally resolve the activity of hundreds of different single-neurons in multiple vertically aligned brain structures. However, while these unprecedented experimental capabilities to study columnar brain properties are a big leap forward in neuroscience, there is the need to spatially distribute electrodes also horizontally. Closely spacing and consistently placing in well-defined geometrical arrangement multiple isolated single-shank probes is methodologically and economically impractical. Here, we present the first high-density CMOS neural probe with multiple shanks integrating thousand’s of closely spaced and simultaneously recording microelectrodes to map neural activity across 2D lattice. Taking advantage from the high-modularity of our electrode-pixels-based SiNAPS technology, we realized a four shanks active dense probe with 256 electrode-pixels/shank and a pitch of 28 *µ*m, for a total of 1024 simultaneously recording channels. The achieved performances allow for full-band, whole-array read-outs at 25 kHz/channel, show a measured input referred noise in the action potential band (300-7000 Hz) of 6.5 ± 2.1*µ*V_*RMS*_, and a power consumption <6 *µ*W/electrode-pixel. Preliminary recordings in awake behaving mice demonstrated the capability of multi-shanks SiNAPS probes to simultaneously record neural activity (both LFPs and spikes) from a brain area >6 mm^2^, spanning cortical, hippocampal and thalamic regions. High-density 2D array enables combining large population unit recording across distributed networks with precise intra- and interlaminar/nuclear mapping of the oscillatory dynamics. These results pave the way to a new generation of high-density and extremely compact multi-shanks CMOS-probes with tunable layouts for electrophysiological mapping of brain activity at the single-neurons resolution.

## 1. Introduction

Here, we present the first high-density CMOS neural probe with multiple shanks integrating thousand’s of closely spaced and simultaneously recording microelectrodes to map neural activity across 2D lattice. Taking advantage from the high-modularity of our electrode-pixels-based SiNAPS technology, we realized a four shanks active dense probe with 256 electrode-pixels/shank and a pitch of 28 *µ*m, for a total of 1024 simultaneously recording channels. The achieved performances allow for full-band, whole-array read-outs at 25 kHz/channel, show a measured input referred noise in the action potential band (300-7000 Hz) of 6.5 ± 2.1*µ*V_*RMS*_, and a power consumption <6 *µ*W/electrode-pixel. Preliminary recordings in awake behaving mice demonstrated the capability of multi-shanks SiNAPS probes to simultaneously record neural activity (both LFPs and spikes) from a brain area >6 mm^2^, spanning cortical, hippocampal and thalamic regions. High-density 2D array enables combining large population unit recording across distributed networks with precise intra- and interlaminar/nuclear mapping of the oscillatory dynamics. These results pave the way to a new generation of high-density and extremely compact multi-shanks CMOS-probes with tunable layouts for electrophysiological mapping of brain activity at the single-neurons resolution. Combining the two facets of multichannel extracellular recordings requires high-resolution, large-scale sensing devices capable of monitoring spiking and local field potentials (LFP) within and across widely distributed circuits (Alivisatos et al. (2013); Buzsáki (2004); Lewis et al. (2015)) and calling for radical scaling of the number of recorded electrodes to yield dense sampling across distributed circuits. Recent studies (Jun et al. (2017b); Raducanu et al. (2016); De Dorigo et al. (2018); Angotzi et al. (2019)) proposed micro-/nano fabricated implantable Complementary Metal-Oxide Semiconductor (CMOS) probes with hundreds to thousands recordings sites within small cross-sectional areas. By integrating into the same silicon substrate electrodes and active electronic circuits for signal amplification and filtering, such implantable CMOS probes can simultaneously record from multiple brain regions (Seymour et al. (2017); Steinmetz et al. (2018)) with an unprecedented spatial resolution. Furthermore, the high 2D spatial resolution of these probes also permits the use of advanced automated sorting algorithms to better isolate putative single-unit activity, by exploiting correlations of individual neuron’s spikes measured by closely and regularly spaced electrode contacts (Hilgen et al. (2017); Jun et al. (2017a); Yger et al. (2018)).

Current CMOS-based probe implementations, however, are limited to linear designs whereby electrodes are arranged along the vertical/axial dimension of implantation, thus being unable to sample anatomical structures that span several millimeters along the horizontal axis and providing mostly serendipitous opportunity for sampling across networks along the axis of the probe. Ability to densely sample extracellular signals for both single neuronal populations and oscillatory dynamics analysis with anatomical resolution requires precise geometrical scaling of the CMOS-based shafts.

In order to address this and many other biological questions related to any type of brain circuits in which vertical and horizontal information needs to be simultaneously sampled, we present here a novel CMOS-probe that exploits the modularity of our SiNAPS (Simultaneous Neural Active Pixel Sensor) technology (see Angotzi et al. (2019)) to integrate simultaneous sampling of 1024 electrode-pixels (electrode-pixel size 26 × 26 *µ*m^2^, pitch 28 *µ*m, electrode pad size 15 × 15 *µ*m^2^) arranged on four distinct shanks (80 *µ*m wide x 5 mm long; incorporating 256 contacts/shank covering 3.6 mm; shank separation 560 *µ*m). Preliminary *in vivo* acute recordings from the brain of behaving head fixed mice demonstrate how broadband neural signals can be acquired from all electrodes simultaneously, with low noise, 6.5±2.1*µ*V_*RMS*_ in the action potential and 8.5 ± 2.7*µ*V_*RMS*_ in the LFP (local field potential) bands, and high temporal resolution (16 kHz sampling frequency).

## 2. Results

### 2.1. SiNAPS 4-shanks probe

The 4-shanks SiNAPS probe solution builds on power efficient analog front-end module that was also used in our single shank device previously described in Angotzi et al. (2019). As illustrated in Figure 1A, a single SiNAPS module comprises 32 electrode-pixel sites, each providing *in situ* amplification and low-pass filtering (Gain = 46 dB, f_*-*3*dB*_ = 4 kHz). These 32 electrode-pixels are time division multiplexed to a variable gain amplifier (up to a 7-fold gain) before off-chip analog to digital conversion (12bit resolution, up to 25 ksamples/s per electrode-pixel). Furthermore, an active feedback circuit shared in a time division multiplexed fashion among the 32 electrode-pixels of the analog frontend periodically and automatically adjusts the operating point of the in-pixel low-noise amplifiers thus correcting for inevitable DC drifts arising at the tissue-electrode electrode-pixel interface and keeping the amplifier within its linear range of operation. This DC offset correction procedure was deemed favorable as it negates the need for additional circuit elements such as RC networks or large feedback capacitors that are subject to mismatch, can degrade input impedance and may lead to signal attenuation.

**Figure 1:**
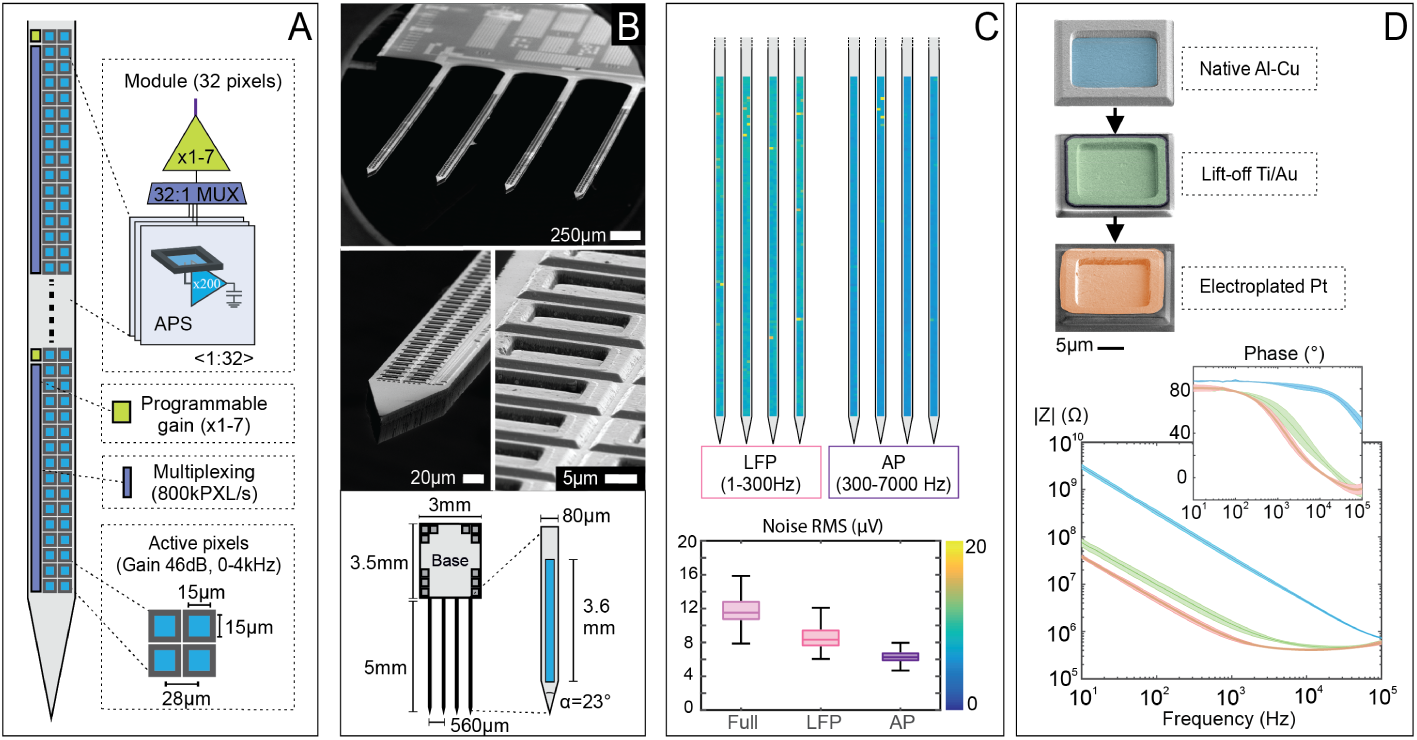
Overview of the multi-shanks SiNAPS technology. **(A)** Architecture and dimensions of the implantable CMOS-probe: multiple modules comprising 32 active electrode-pixels that are time division multiplexed to a variable gain amplifier are arranged over a 80 *µ*m wide shank for a total of 256 electrode-pixels per shank (pitch 28 *µ*m, electrode pad size 15 × 15 *µ*m^2^). **(B)** Scanning Electron Micrographs showing a structured 4-shanks SiNAPS at different magnification levels. Shanks are 5mm long, with active sensing of 3.6 mm in length, and 560 *µ*m shanks separation. **(C)** Noise performances in saline of a 4-shanks SiNAPS probe in the Local Field Potential (LFP, 1-300 Hz) and the Action Potential (AP, 300-7000 Hz) bandwidth, respectively 8.5±2.7*µ*V_*RMS*_ and 6.5±2.1*µ*V_*RMS*_. Figure also reports the broadband (1-7000 Hz) noise distribution across channels (11.9 ± 3.7*µ*V_*RMS*_). **(D)** False-coloured SEM images and corresponding Electrochemical Impedance Spectroscopy measurements at different steps of active pixels material improvements. The native Al-Cu alloy issued from the foundry reveals a mean electrode-pixel impedance of 35.9 ± 0.4 MΩ @1kHz. Evaporation and lift-off of Ti/Au 10/100 nm on top of the Al-Cu electrode-pixels reduce this value to 1.4±0.3 MΩ, while further Pt electroplating reaches a mean electrode-pixel impedance of 0.7±0.1 MΩ (N=3 probes×1024 electrodes).

As illustrated in Figure 1A-B, with the aim of minimizing damages of brain tissue during implantation, in this current implementation the 32 electrode-pixels comprising the analog front-end module are arranged with a 28 *µ*m pitch in two rather than three columns as for the single shank SiNAPS probe, thus reducing the cross-sectional width of the shanks to 80 *µ*m. Each of the four shanks integrates eight of such modules, and achieves a total number of 256 active recording sites over an active length of 3.58 mm. Finally, the inter-shank separation of about 560 *µ*m was specifically set to target large portions of the hippocampus in rodents brain. The overall CMOS area needed to fabricate each multi-shanks SiNAPS probe (base + shanks) is <12 mm^2^, with an electrode-pixel density close to 90 pixels/mm^2^. This confirms one of the beneficial features of the SiNAPS technology to be at least one order of magnitude denser than other CMOS neural probes in terms of simultaneously recording channels per silicon area.

By optimizing on-chip circuits with respect to noise, area and power consumption, we achieved total electrode-pixel size of 26 *µ*m ×26 *µ*m, a uniform low noise in the action potential (AP, 0.3-5 kHz) and local field potential (LFP, 0 – 1 kHz) range accounting respectively for about 6.5±2.1 *µ*V and 8.5±2.7 *µ*V root-mean square (rms), while consuming only 6 *µ*W of power per electrode-pixel (see Figure 1C). An NMOS transistor working as electronic switch is integrated in each electrode-pixel, permitting to connect the metal electrode to a common metal line that is routed off chip. This ultimately provides an electrical path to the metal electrodes that can be used for either circuit characterization or for electroplating the native CMOS Al-Cu alloy at the electrode sites with more bio-compatible and lower impedance materials (see section Materials & Methods for details). Specifically, thermally evaporated gold lift-off deposition on top of the native Al-CU and further platinum electroplating reduced the native 35.9±0.4 MΩ electrode impedance (Al-Cu) down to 1.4±0.3 MΩ (Au) and 0.7±0.1 MΩ (Pt) respectively (Figure 1D).

### 2.2. Acute in vivo recordings

To assess multi-shanks SiNAPS probe performance we conducted acute recordings on awake and anaesthetized head fixed and anaesthetized rodents. We simultaneously acquired broadband signals (1-7500 Hz) from different brain structures, including parietal cortex, hippocampus and the underlying thalamic nuclei, towards a widespread high spatiotemporal resolution bioelectrical mapping of large planar brain regions (Figure 2A). From the AP band pass filtered neural data (300-5000 Hz) we isolated single units using the Kilosort2 software (Pachitariu et al. (2016)) followed by manual curation using the Phy graphical user interface (https://github.com/cortex-lab/phy). Following this procedure we were able to sort hundreds of well-isolated neurons distributed across the three major brain structures targeted in our experiments (cortex, hippocampus and thalamus). Figure 2B. reports results from one representative experimental trial.

**Figure 2:**
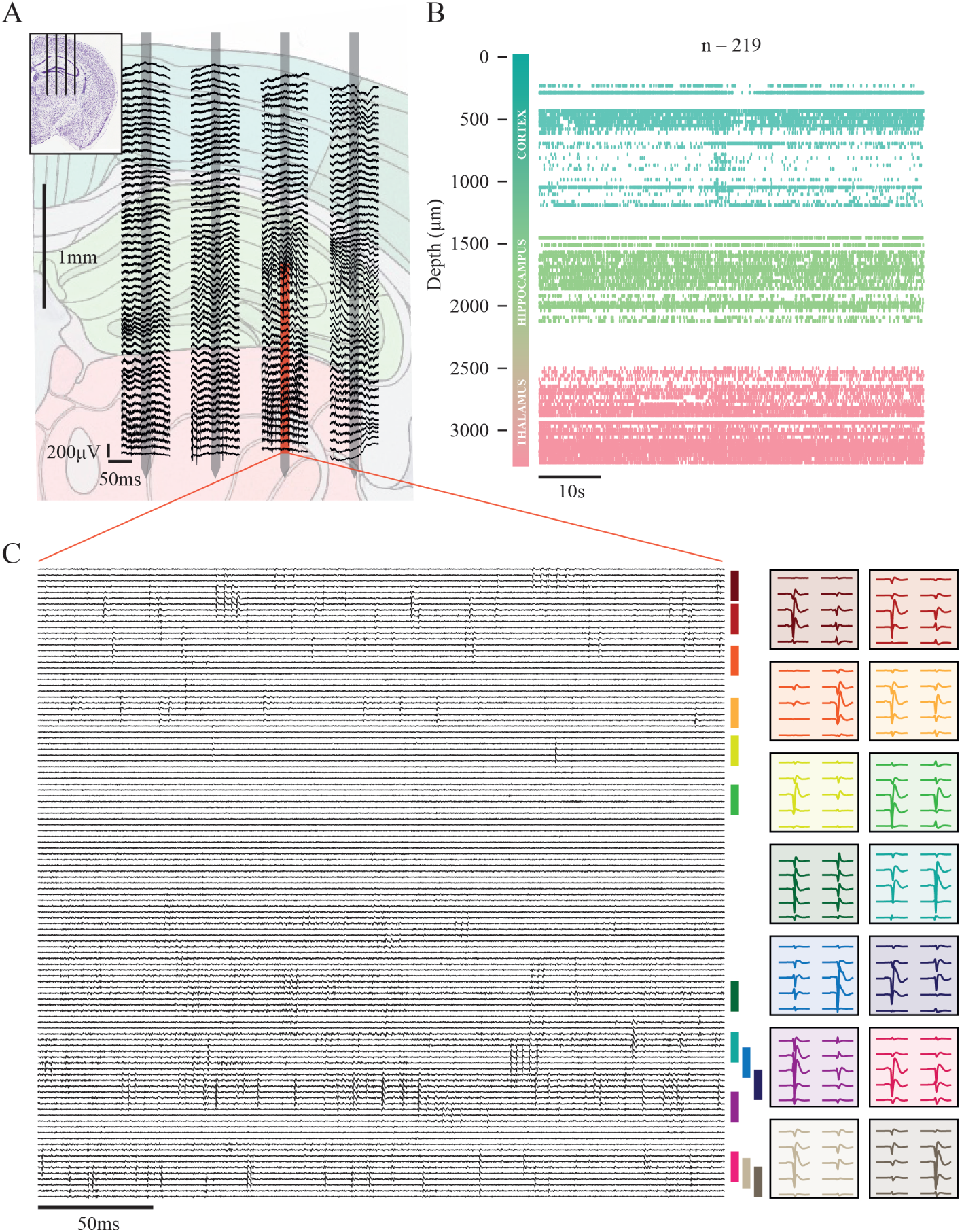
In vivo recording with multi-shanks SiNAPS probe. **(A)** Approximate probe position during experiments superimposed to Allen Mouse Brain Atlas (−2 mm AP from bregma) with, for each shank, 100 ms of the broadband neural signals collected from a representative subset of the 1024 available channels. **(B)** Raster plot of the single unit (n = 219) activity recorded from the entire probe. Each color represents the macro structure from which a specific neuron has been recorded.**(C)** Example of band pass filtered (300-5000 Hz) neural data collected from a portion of one of the shank (red area composed by 108 neighbouring consecutive channels) of the probe depicted in (A). On the right side of the picture are reported the waveforms of a subset of the units recorded from those channels. Color coding represents the position in the raw data from which the units were acquired (their spatial extent is represented by the colored rectangle on the right side of the neural traces).

Low-noise and high-count densely packed recording electrodes allowed, as for single shank high-density probes, to record a large variety of neurons, that differ in amplitudes and shapes, and to isolate multiple cells in small local areas. The capability to isolate single neuron waveshapes in wide brain portion also allows to dissect the coordinate functions of different cell types across distributed circuits (Sirota et al. (2008),Fernández-Ruiz et al. (2017)). For example in a portion of 108 consecutive channels in one of the four available shanks, we were able to record the activity of more than one hundred of different neurons (for clarity only a subset of these units is reported in Figure 2C).

It is worth mentioning that While in some cases we were unable to lower the full 256 channels of each shank into the brain due to space constraints, we noticed that there was useful information to be obtained at the boundary of electrodes in and out of the brain. Specifically we were able to extract information about the curvature of the brain that could be used post hoc to localize the electrode positions. Coupled with the continuous acquisitions from all 1024 electrode-pixels, one can infer the relative position of each shank with respect to the cortical surface (at the electrodes pitch resolution) based on the recorded signals (see supplementary Figure S1).

## 3. Materials and methods

### 3.1. CMOS post-processing

As for the single shank device, our current 4-shanks implementation was designed and fabricated in a standard 180 nm Complementary Metal-Oxide Semiconductor (CMOS) technology. Devices were delivered as single dies with size of (W ×L×T = 3 mm×8.5 mm×250 *µ*m) and were post processed in our clean room to make them compatible for brain implantation (final size of each shank W×L×T = 80 *µ*m×5 mm×30 *µ*m). The process flow involves a two-step lithography: a lift-off of Ti/Au on top of the native Al-Cu electrode-pixels to improve the neural interfaces with a noble metal of lower impedance, then the standard process flow established previously for shaping and thinning (Angotzi et al. (2019)). Briefly, multiple single chips are fixed onto a single glass slide using water-dissoluble glue, and 80% of the steps in the process can be performed on several dies simultaneously. Both lithographies are performed using spray-coating of photoresist, which enables a more conformable coating on top of these singles dies. For the lift-off, 10/100 nm of Ti/Au are thermally evaporated in the shape of 20×20 *µ*m squares on top of the native Al-Cu electrode-pixels, providing a uniform, electrochemically-stable metal layer to interface with the neurons. The second lithography involves chemical etching of a 400 nm sputtered Cr layer to define the shanks width and length, then a Bosch process to shape these shanks. Finally, the dies are flipped upside down and the Bosch is repeated on the backside until reaching desired shank thickness. The pad area, on the other hand, remains intact. Finally, the resulting structured SiNAPS probes are wire-bonded on printed circuit boards (PCB).

### 3.2. SiNAPS acquisition system

As previously described in Angotzi et al. (2019) the back-end module (currently capable of simultaneous, real-time acquisition of 4096 channels with a 25 kHz sampling rate) comprises an interface board, an acquisition unit, and a data-acquisition software running on a PC equipped with a frame grabber. The interface board connects the headstage mounting the SiNAPS probe to the acquisition unit and integrates a bank of 32 analog to digital converters, one for each of the 32 electrode-pixels module of the SiNAPS probe (MAX11105, Maxim Integrated, USA) with 12-bit resolution each. The acquisition unit is designed on an Opal Kelly ZEM4310 integration module based on an Altera Cyclone IV FPGA. The unit implements a controller for the ADCs, an input memory buffer, a bank of programmable band-pass digital filters, a bidirectional interface for communication with the CMOS-probe, and a Cameralink interface for real-time, high-rate data transmission to a PC. Finally, the PC runs a data acquisition software (BrainWave, 3BrainAG, Switzerland) for data visualization and storage which is currently limited to a maximum acquisition rate of about 16 ksample/s. Because of such limitation, data presented in this work was acquired with a sampling rate of 15500 Hz.

### 3.3. Surgical procedures and acute in vivo recordings

Three adult male mice were prepared for these experiments but only data from a single mouse are shown in this work. Mice were first habituated to human handling after which surgical procedures were administered. Anaesthetised was induced using a midazolam, medetomiding and fentanyl cocktail. Mice were then headfixed within a stereotaxic frame and oxygenated air was supplied at 1 l/min throughout the surgical procedures. Anaesthesia maintenance was achieved with 0.5-1% isoflurane which was administered through a vaporiser. After rostral skin incision and clearence of the skull from any connective tissue, several screws were implanted around the location of the craniotomy destined to serve as ground and reference screws. Two of these were found on the occipital bone above the cerebellum, while a third was implanted very close to the point of insertion of the SiNAPS4 probe (medial shank coordinates AP:-2 mm, ML: +0.9 mm). We observed no differences in signal quality when using any of these reference screws. To enable a stable head fixation of the animal during behavioral recordings, a custom designed head post was implanted onto the posterior bone plate. An additional wire was inserted into the olfactory sensory epithelium allowing recordings of respiration rhythm. All screws and headpost were secured onto the skull using dental acrylic cement. Finally a cranial window was drilled above the dorsal hippocampus, the dura was carefully retracted and the craniotomy was sealed with a two component artificial dura that was easily penetrable using multishank probes. A further layer of a silicon elastomer (quick seal) was added above but not in contanct with the artificial dura that protected the craniotomy and cortex from any mechanical stress. After a week of post operative recovery, mice were water deprived and trained to run on a running disk for water reward. The animals’running speed controlled in a closed loop fashion the speed of a carousel environment that rotated infront of the animal. This carousel environment allowed for analysis of virtual spatial tuning of single cells to the carousel environment. On recording days, the SiNAPS probe was mounted on a micromanipulator, aligned to the craniotomy and lowered down to 3.4 mm from the brain surface (insertion speed 0.2 *µ*m/s). After 15 minutes of resting, in which the animal calmed down in the complete darkness, we acquired from the multi-shanks SiNAPS probe broadband neural activity from cortical, hippocampal and thalamic structures. After recordings, the probe was extracted from the mouse brain and rinsed in distilled water to be reused in the next experimental sessions. All procedures were performed in accordance with the European Communities Directive 2010/63/EC and the German Law for Protection of Animals (TierSchG), and were approved by local authorities (ROB-55.2-2532.Vet 02-16-170). All efforts were made to minimize the number of animals used and the incurred discomfort.

### 3.4. Electrical and Electrochemical characterization

Devices were tested with respect to their electrical and electrochemical performances prior to *in vivo* acute recordings using two methods. In the first method, the noise contribution of the electronic circuits is estimated by using the switch that is integrated in each electrode-pixel allowing to connect all the electrodes to a common ground. The second method involves noise contribution measurements in wet conditions by dipping the shanks in a beaker containing saline solution (NaCl 0.9%) that is forced to ground. Data collected at the output of the whole signal acquisition chain under these two different configurations is used to compute the power spectral density and, thus, the noise contribution within specific frequency bands.

In addition, electrochemical impedance spectroscopy (EIS) is performed in NaCl 0.9% solution at ten frequencies per decade over the range 101–105 Hz (PG-STAT204, Metrohm Autolab, Switzerland). The impedance value for the single electrode can be estimated as 1024× larger than the one measured with all the 1024 sensing electrodes connected in parallel.

## 4. Discussions

A number of different approaches have been recently proposed to target single neuron recordings from wide distributed brain circuits wit sub-millisecond temporal resolutions. High channel count solutions based on the use of thin, flexible multi-electrode polymer probes were recently proposed (Chung et al. (2019), Musket al. (2019)), but their efficacy remains to be demonstrated. These approaches are likely to suffer from several drawbacks, including electromagnetic interference (EMI), signal attenuation due to parasitic capacitances, and crosstalk between adjacent channels due to the densely packed metal lines. Furthermore, such thin polymer probes are not stiff enough to be directly inserted into the brain and their use requires additional solutions and steps for their routine implementation.

Precise geometrical tissue sampling by such probes is also not currently feasible. Conversely, active implantable probes based on CMOS technology surpass all the above mentioned limitations. These include Neuropixels (Jun et al. (2017b)), Neuroseeker (Raducanu et al. (2016)) and, our own SiNAPS solution (Angotzi et al. (2019)). Among these options, SiNAPS probes are the only devices that overcome the major scaling bottleneck caused by the spatial limits of analog front-ends (Seymour et al. (2017)), thus allowing on-probe multiplexing of thousands of electrode signals on a few output lines. This approach has tremendous scaling potential: by increasing the number of recording electrodes while minimizing overall probe size, particularly the base area, SiNAPS is also a very promising solution for chronic experiments in animals, especially small animals like mice. Moreover, because CMOS fabrication costs are directly proportional to the circuit area, the raw fabrication costs of SiNAPS probes can be much lower than other available high-density CMOS probes, favoring SiNAPS’ widespread dissemination in the neuroscience community.

In detail, like Neuropixels and Neuroseeker, SiNAPS use CMOS technology to integrate similarly dense arrays of electrodes and read-out circuits but with radically different concepts leading to distinct advantages. Specifically, SiNAPS probes implement the Active Pixel Sensor (APS) concept originally developed for image sensors (Fossum (1997)), whereby active circuits for signal amplification, low-pass filtering and read-out are located directly underneath each electrode-pixel (Berdondini et al. (2001)). Notably, this permits integrating an equal number of front-ends and electrodes as required for simultaneous recordings from the entire electrode array in contrast with other technologies where only a subset of channels can be simultaneously sampled. Furthermore, the unique underlying engineering and modular design of our SiNAPs probes allows to rapidly augment the probes’ capabilities well beyond those of currently available commercial probes. These augmentations include scaling up the number of (fully multiplexed) electrode contacts far beyond that of any competing probe with user-desired shaft configurations and geometries. As demonstrated in this work, current SiNAPS probes can record concurrently from 1024 electrode-pixels distributed across four equally spaced shanks at 16 ksamples/s per electrode. This unprecedented spatiotemporal sampling of large brain areas (more than 6 mm^2^ with ∼ 170 recording electrode-pixels per square millimeter) can permit a much finer description and comprehension of neural dynamics across different structures while overcoming most of the drawbacks intrinsically linked to passive neural probes such as large footprint, power consumption and wiring (Berenyi et al. (2014), Shobe et al. (2015)). Preliminary data presented here confirm and advance SiNAPS capabilities previously demonstrated in (Angotzi et al. (2019)) with respect to noise performances and neural signal quality. Furthermore, it also show, for the first time, the possibility to simultaneously sample large brain areas in the bidimensional space with fine spatial resolution. We strongly believe that our multi-shanks technology will significantly contribute in advancing the tools that have been originally developed over the last few years for the post-processing and interpretation of the rich spatio-temporal content of data collected from high-density single shank neural probes (Jun et al. (2017a), Hilgen et al. (2017), Pachitariu et al. (2016), Rossant et al. (2016), Yger et al. (2018)). Ultimately, this will lead to a better understanding of the implementation and execution of distributed brain functions.

## Author contributions

L.B. and G.N.A. coordinated the SiNAPS project. L.B., G.N.A. and F.B. designed the study and wrote the manuscript. G.N.A. developed the multi-shanks SiNAPS probes and the whole back-end acquisition system. A.L. was responsible for the CMOS post-processing and electrochemical characterization. A.S., N.P. and G.S. designed and performed the animal experiments, analyzed data and contributed to the manuscript. F.B. and S.Z. contributed to the animal experiments and analyzed the data. All co-authors reviewed the paper.

## Acknowledgements

This project has received funding from the European Union’s Horizon 2020 research and innovation programme under the Marie Sklodowska-Curie grant agreement No 746778.

---

**Figure S1:**
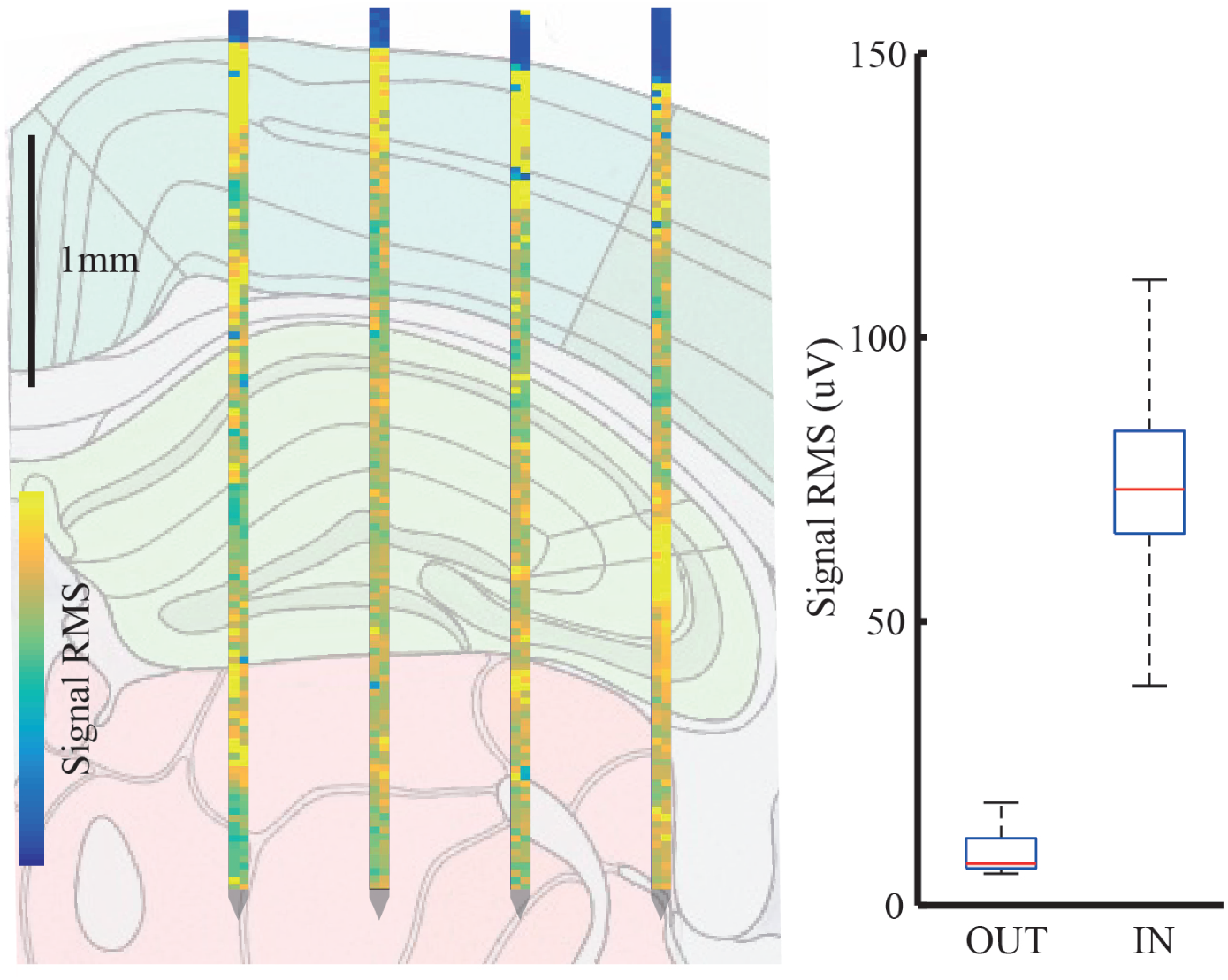
On-line localization of SiNAPS probe. False color map in which each electrode-pixel represents the signal RMS measured from an electrode in a 5 seconds window during an *in vivo* recording session. Overlaid Brain Atlas image reveals the close match between the RMS variation edge (from like-blue to like-yellow pixels) and the cortical brain surface. On the right side the boxplot reports the RMS values distribution of the electrode-pixels outside the brain then respect to the one inside. False color map has been superimposed to the atlas image through the depth of the most medial shank displayed by the micromanipulator during the experimental session.

## Bibliography

Alivisatos, A. P., Andrews, A. M., Boyden, E. S., Chun, M., Church, G. M., Deisseroth, K., Donoghue, J. P., Fraser, S. E., Lippincott-Schwartz, J., Looger, L. L., Masmanidis, S., McEuen, P. L., Nurmikko, A. V., Park, H., Peterka, D. S., Reid, C., Roukes, M. L., Scherer, A., Schnitzer, M., Sejnowski, T. J., Shepard, K. L., Tsao, D., Turrigiano, G., Weiss, P. S., Xu, C., Yuste, R., Zhuang, X., 2013. Nanotools for neuroscience and brain activity mapping. ACS Nano 7 (3), 1850–1866.

Angotzi, G. N., Boi, F., Lecomte, A., Miele, E., Malerba, M., Zucca, S., Casile, A., Berdondini, L., feb 2019. SiNAPS: An implantable active pixel sensor CMOS-probe for simultaneous large-scale neural recordings. Biosensors and Bioelectronics 126, 355–364.

Berdondini, L., Overstolz, T., de Rooij, N., Koudelka-Hep, M., Wany, M., Seitz, P., 2001. High-density microelectrode arrays for electrophysiological activity imaging of neuronal networks. In: ICECS 2001. 8th IEEE International Conference on Electronics, Circuits and Systems (Cat. No.01EX483). Vol. 3. IEEE, pp. 1239–1242.

Berenyi, A., Somogyvari, Z., Nagy, A. J., Roux, L., Long, J. D., Fujisawa, S., Stark, E., Leonardo, A., Harris, T. D., Buzsaki, G., 2014. Large-scale, high-density (up to 512 channels) recording of local circuits in behaving animals. Journal of Neurophysiology 111 (5), 1132–1149.

Buzsáki, G., may 2004. Large-scale recording of neuronal ensembles. Nature Neuroscience 7 (5), 446–451.

Chung, J. E., Joo, H. R., Fan, J. L., Liu, D. F., Barnett, A. H., Chen, S., Geaghan-Breiner, C., Karlsson, M. P., Karlsson, M., Lee, K. Y., et al., 2019. High-density, long-lasting, and multi-region electrophysiological recordings using polymer electrode arrays. Neuron 101 (1), 21–31.

De Dorigo, D., Moranz, C., Graf, H., Marx, M., Wendler, D., Shui, B., Herbawi, A. S., Kuhl, M., Ruther, P., Paul, O., et al., 2018. Fully immersible subcortical neural probes with modular architecture and a delta-sigma adc integrated under each electrode for parallel readout of 144 recording sites. IEEE Journal of Solid-State Circuits 53 (11), 3111–3125.

Fernández-Ruiz, A., Oliva, A., Nagy, G. A., Maurer, A. P., Berényi, A., Buzsáki, G., 2017. Entorhinal-ca3 dual-input control of spike timing in the hippocampus by theta-gamma coupling. Neuron 93 (5), 1213–1226.

Fossum, E., 1997. CMOS image sensors: electronic camera-on-a-chip. IEEE Transactions on Electron Devices 44 (10), 1689–1698.

Hilgen, G., Sorbaro, M., Pirmoradian, S., Zanacchi, F. C., Sernagor, E., Hennig, M. H., Hilgen, G., Sorbaro, M., Pirmoradian, S., Muthmann, J.-o., Kepiro, I. E., Sona, D., Zanacchi, F. C., Sernagor, E., Hennig, M. H., 2017. Unsupervised Spike Sorting for Large-Scale, High-Density Multi-electrode Arrays Resource Unsupervised Spike Sorting for Large-Scale, High-Density Multielectrode Arrays. CellReports 18 (10), 2521–2532.

Jun, J. J., Mitelut, C., Lai, C., Gratiy, S., Anastassiou, C., Harris, T. D., 2017a. Real-time spike sorting platform for high-density extracellular probes with ground-truth validation and drift correction. bioRxiv, 101030.

Jun, J. J., Steinmetz, N. A., Siegle, J. H., Denman, D. J., Bauza, M., Barbarits, B., Lee, A. K., Anastassiou, C. A., Andrei, A., Aydin, Ç., Barbic, M., Blanche, T. J., Bonin, V., Couto, J. J., Dutta, B., Gratiy, S. L., Gutnisky, D. A., Häusser, M., Karsh, B., Ledochowitsch, P., Lopez, C. M., Mitelut, C., Musa, S., Okun, M., Pachitariu, M., Putzeys, J., Rich, P. D., Rossant, C., Sun, W. L., Svoboda, K., Carandini, M., Harris, K. D., Koch, C., O’Keefe, J., Harris, T. D., Aydin, C., Barbic, M., Blanche, T. J., Bonin, V., Couto, J. J., Dutta, B., Gratiy, S. L., Gutnisky, D. A., Hausser, M., Karsh, B., Ledochowitsch, P., Lopez, C. M., Mitelut, C., Musa, S., Okun, M., Pachitariu, M., Putzeys, J., Rich, P. D., Rossant, C., Sun, W. L., Svoboda, K., Carandini, M., Harris, K. D., Koch, C., O’Keefe, J., Harris, T. D., 2017b. Fully Integrated Silicon Probes for High-Density Recording of Neural Activity. Nature in press (7679), 232–236.

Lewis, C. M., Bosman, C. A., Fries, P., 2015. Recording of brain activity across spatial scales.

Musk, E., et al., 2019. An integrated brain-machine interface platform with thousands of channels. BioRxiv, 703801.

Pachitariu, M., Steinmetz, N. A., Kadir, S. N., Carandini, M., Harris, K. D., 2016. Fast and accurate spike sorting of high-channel count probes with kilosort. In: Advances in Neural Information Processing Systems. pp. 4448–4456.

Raducanu, B. C., Yazicioglu, R. F., Lopez, C. M., Ballini, M., Putzeys, J., Wang, S., Andrei, A., Welkenhuysen, M., Van Helleputte, N., Musa, S., Puers, R., Kloosterman, F., Van Hoof, C., Mitra, S., 2016. Time multiplexed active neural probe with 678 parallel recording sites. European Solid-State Device Research Conference 2016-Octob (10), 385–388.

Rossant, C., Kadir, S. N., Goodman, D. F., Schulman, J., Hunter, M. L., Saleem, A. B., Grosmark, A., Belluscio, M., Denfield, G. H., Ecker, A. S., et al., 2016. Spike sorting for large, dense electrode arrays. Nature neuroscience 19 (4), 634.

Seymour, J. P., Wu, F., Wise, K. D., Yoon, E., 2017. State-of-the-art MEMS and microsystem tools for brain research. Microsystems & Nanoengineering 3, 16066.

Shobe, J. L., Claar, L. D., Parhami, S., Bakhurin, K. I., Masmanidis, S. C., 2015. Brain activity mapping at multiple scales with silicon microprobes containing 1,024 electrodes. Journal of neurophysiology 114 (3), 2043–2052.

Sirota, A., Montgomery, S., Fujisawa, S., Isomura, Y., Zugaro, M., Buzsáki, G., 2008. Entrainment of neocortical neurons and gamma oscillations by the hippocampal theta rhythm. Neuron 60 (4), 683–697.

Steinmetz, N. A., Koch, C., Harris, K. D., Carandini, M., 2018. Challenges and opportunities for large-scale electrophysiology with Neuropixels probes. Current Opinion in Neurobiology 50, 92–100.

Yger, P., Spampinato, G. L., Esposito, E., Lefebvre, B., Deny, S., Gardella, C., Stimberg, M., Jetter, F., Zeck, G., Picaud, S., Duebel, J., Marre, O., 2018. A spike sorting toolbox for up to thousands of electrodes validated with ground truth recordings in vitro and in vivo. eLife 7, e34518.

